# Rate of osmotic pressure change in drying saliva microdroplets drives inactivation of surrogate respiratory bacteria

**DOI:** 10.64898/2026.05.19.726210

**Authors:** Taylor Medina, Beiping Luo, Thomas Peter, Htet Kyi Wynn, Tamar Kohna

## Abstract

Airborne transmission of respiratory pathogens depends on their ability to remain viable in drying respiratory droplets, yet the physicochemical drivers of bacterial inactivation during droplet evaporation remain poorly quantified. This study combines controlled droplet experiments with physicochemical modeling to investigate how osmotic pressure dynamics influence bacterial survival. Using *Escherichia coli* and *Staphylococcus epidermidis* as Gram-negative and Gram-positive surrogates, respectively, we measured viability loss in artificial saliva droplets dried at multiple relative humidities and reconstructed the time-resolved osmotic pressure using the Respiratory Aerosol Model (ResAM). Both organisms remained stable while droplets were liquid but lost viability following efflorescence, when rapid solute concentration changes produced sharp osmotic pressure increases. The extent of inactivation scales log-linearly with the rate of osmotic pressure change around efflorescence: *E. coli* decays faster than *S. epidermidis*, and relationships derived in artificial saliva predict survival in independent phosphate-buffered saline experiments. A more rapid drop in humidity led to more severe osmotic shocks and greater inactivation. These results identify the rate of osmotic pressure change during efflorescence as a quantitative, medium-independent predictor of bacterial survival in drying respiratory droplets.

**Importance:** Airborne infection risk depends on how long microorganisms remain viable in respiratory particles after exhalation, yet the physical mechanisms controlling bacterial survival during droplet drying are not well defined. Evaporation of respiratory droplets concentrates salts and can impose sudden and extreme osmotic stress on microbes, but this process has been difficult to quantify because osmotic pressure cannot be measured directly inside microscopic droplets. Integration of droplet experiments with a physicochemical aerosol model shows that bacterial inactivation is governed primarily by the rate of osmotic pressure increase during droplet efflorescence rather than by static values of humidity or solute concentration alone.

This mechanism explains why rapid drying may produce strong inactivation.

## Introduction

Respiratory pathogens, including viruses, bacteria, and fungi, are transmitted by the airborne route and exhibit significant variation in their airborne stability. Prominent examples such as influenza viruses, Mycobacterium tuberculosis, and SARS-CoV-2 highlight this diversity and their potential for widespread transmission and outbreaks^1^. These pathogens are transmitted through infectious respiratory particles (IRPs), which include both small respiratory aerosol particles and larger droplets^2^. Successful airborne transmission critically depends on a pathogen’s ability to maintain infectivity in IRPs. Yet, the stability of pathogens in IRPs is only partially understood.

A known driver of pathogen decay in IRPs is the changing salt concentration^3,4^. After IRPs are exhaled, they are subject to evaporation, which is dependent on the ambient relative humidity (RH). Efflorescence occurs when ambient relative humidity falls below the efflorescence relative humidity (ERH), causing dissolved salts to crystallize^5^. However, the presence of a nucleating agent, such as insoluble particles or certain biological materials, can promote efflorescence, meaning that the ERH is then substantially higher^6^. During the drying process, salt concentrations can reach high supersaturations, but the concentration drops back to saturation levels once efflorescence takes place.^6^

Although the specific response to changing salt concentrations in IRPs varies between pathogen types, bacterial inactivation can, at least in part, be attributed to hyperosmotic shock experienced just before droplet efflorescence^7^. This shock arises from a rapid change in solute concentrations in the surrounding solution to levels that exceed those of the intracellular environment. As evaporation proceeds, the increasing solute concentration reduces the water activity (*a*_*w*_), a dimensionless quantity relative to pure water that reflects the chemical potential of water^8^. This leads to a rise in the osmotic pressure (*Π*), a thermodynamic measure of a solution’s tendency to draw in water relative to pure water, proportional to *-ln(a*_*w*_) ^8,9^. For a bacterium within the droplet, this defines the extracellular osmotic environment (*Π*_*droplet*_). Increases in external osmotic pressure drive water efflux from the cytoplasm, resulting in a reduction in turgor pressure (*P*_*turgor*_), the mechanical force generated by the difference between the intracellular (*Π*_*bacteria*_) and extracellular osmotic pressures (*P*_*turgor*_ *= Π*_*bacteria*_ *– Π*_*droplet*_) ^10,11^. Turgor pressure is essential for maintaining cellular shape and serves as a signal for regulating the expression of osmoregulatory genes ^11^. Rapid increases in *Π*_*droplet*_ can therefore cause a sharp decline in *P*_*turgor*_ before osmoadaptive mechanisms can restore intracellular osmotic balance. This transient loss of turgor leads to membrane deformation, mechanical stress on the cell envelope, and ultimately loss of viability, thereby linking evaporation dynamics to pathogen inactivation in IRPs.

*Escherichia coli* and *Staphylococcus epidermidis* represent Gram-negative and Gram-positive bacteria with distinct ecological niches and physiological adaptations. *E. coli* is primarily associated with aqueous environments such as the gastrointestinal tract, whereas *S. epidermidis* is adapted to the human skin, where it is routinely exposed to desiccation and osmotic stress. These differences are reflected in their osmotic stress responses. *E. coli* and other Gram-negative bacteria maintain a turgor pressure of about 2–4 atm through molecular osmoadaptation mechanisms and can tolerate transient osmotic pressure increases up to 10– 20 atm^12^. However, viability depends not only on the magnitude of osmotic pressure increase, but also on the rate of increase, as rapid increases in *Π*_*droplet*_ produce abrupt reductions in *P*_*turgo*r_ that the bacterium cannot compensate for. *E. coli* can only fully recover from shocks up to 22 atm within 10-15 minutes ^8,13^. While the initial response to hyperosmotic stress is initiated within 1–2 minutes, full osmoadaptive gene expression requires approximately one hour, depending on physiological state and media composition^14^. The long response time renders Gram-negative bacteria generally intolerant of rapid osmotic shocks seen during efflorescence.

In contrast, Gram-positive bacteria generally exhibit higher resistance to osmotic and desiccation stress^15^. This resilience is attributed to their thicker peptidoglycan layer, which provides additional mechanical strength and allows the cell wall to sustain higher turgor pressures, often exceeding 20 atm^16^. Elevated turgor pressure, together with the rigid peptidoglycan matrix, buffers the cell against external osmotic shock by reducing the relative change in transmembrane pressure and limiting membrane deformation. This mechanical stabilization mitigates cell lysis and structural damage under high osmotic conditions. As a result, Gram-positive bacteria typically show slower inactivation rates under drying conditions compared to Gram-negative species, which are more vulnerable to rapid water loss and membrane tension spikes^10^.

Despite the recognized importance of osmotic pressure as a driver of bacterial inactivation in IRPs^17,18,19^, there is currently no quantitative approach to describe the dependence of inactivation kinetics on osmotic pressure within these particles. This is partly due to the fact that direct internal measurements of osmotic pressure are not feasible. This study addresses this fundamental gap by combining empirical data on bacterial inactivation with a physicochemical droplet model to estimate osmotic pressure during drying. Specifically, this study tests the hypothesis that bacterial inactivation in drying IRPs is governed primarily by the rate of osmotic pressure increase just before efflorescence. *E. coli* (a Gram-negative bacterium) and *S. epidermidis* (a Gram-positive bacterium) are used to represent different classes of bacteria with varied structural, physiological, and environmental susceptibilities. These organisms are commonly used as non-pathogenic surrogate organisms to avoid working with pathogens, which require increased biosafety containment. The study first quantifies the inactivation kinetics of *E. coli* and *S. epidermidis* in 1 μL deposited artificial saliva droplets under controlled RH conditions and then employs the physicochemical model ResAM to model the osmotic pressure over time inside the droplet throughout the drying process^4^. This integrated approach allows the establishment of a quantitative correlation between modeled change in osmotic pressure just before efflorescence and microbial viability in drying respiratory droplets, providing mechanistic insight into bacterial survival.

## Results

### Evolution of Bacterial Viability and Osmotic Pressure in Drying Saliva Droplets

To determine how relative humidity (RH) influences bacterial survival during droplet drying, *E. coli* and *S. epidermidis* suspended in 1 μL artificial saliva droplets were deposited on hydrophobic-coated 96 well plates and were dried at low (30%), medium (50%), and high (70%) RH. Bacterial viability in the droplets was monitored for 60 minutes at 30% RH for *E*. coli and for 120 minutes for the rest of the experiments. Parallel simulations using ResAM modeled the evolution of *Π*_droplet_ based on droplet composition, RH and observed droplet radius.

Figure 1 shows bacterial inactivation curves together with the modeled osmotic pressure. The osmotic pressures reported here are referenced to pure water and capture the magnitude and rate of increase in the extracellular osmotic environment, thereby representing the osmotic shock experienced by the bacterium.

**Figure 1.**
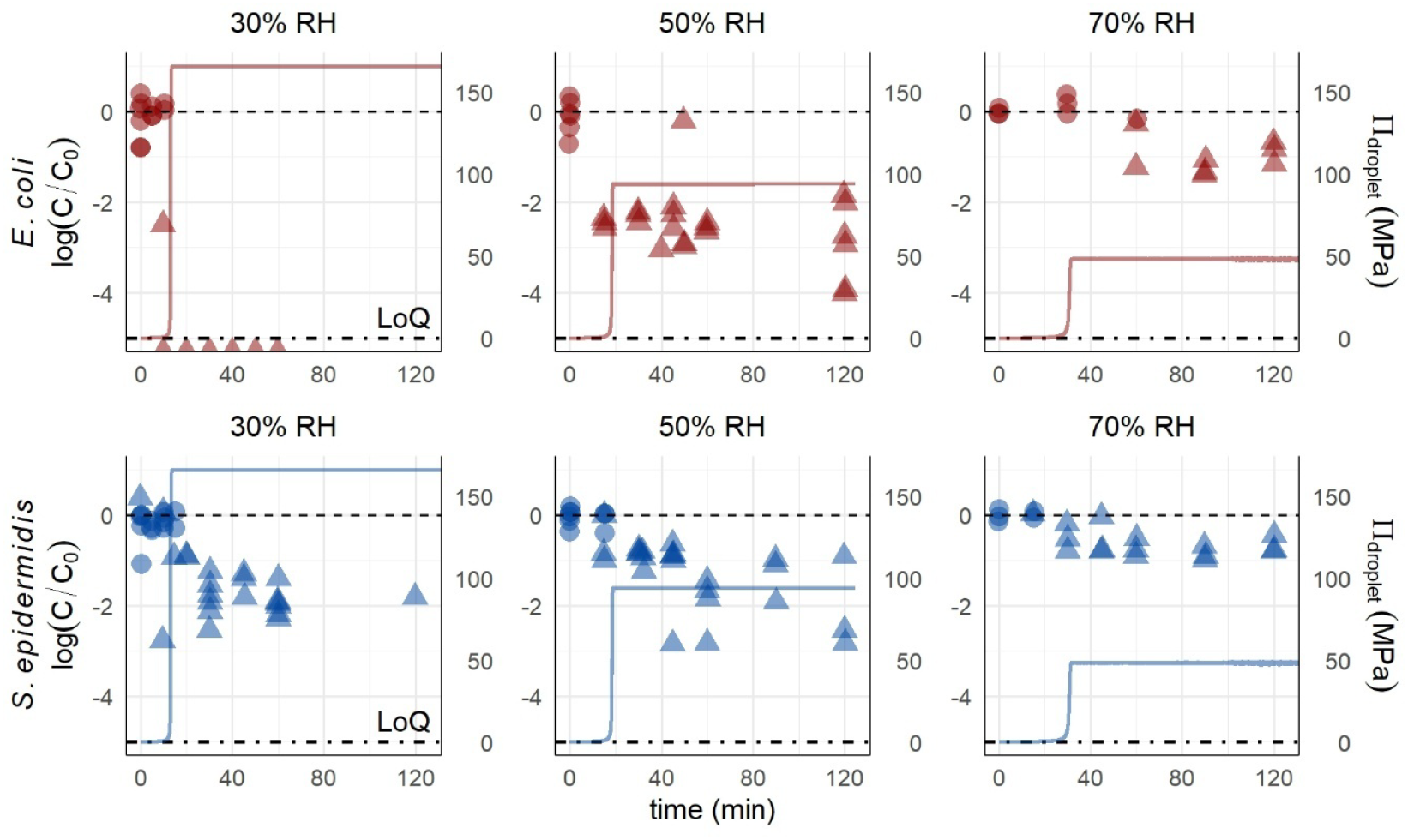
Inactivation of *E. coli* (red) and *S epidermidis* (blue) in 1 μL artificial saliva droplets dried at 30%, 50%, and 70% relative humidity (RH) with modeled osmotic pressure (*Π*_droplet_). The left axis represents the log_10_ reduction of the bacteria, while the right axis shows the modeled osmotic pressure *Π*_droplet_ (MPa) shown as a solid line. Circles denote liquid droplets, and triangles represent effloresced droplets. The starting titers of *E. coli* and *S. epidermidis* were both 10^8^ CFU/mL.

During the initial drying phase, osmotic pressure changed only slightly as water evaporated (Figure 1). As solutes approached supersaturation near efflorescence, osmotic pressure increased sharply, followed by a brief dip associated with crystallization, and then a final rise to the equilibrium osmotic pressure determined by RH. The modeled dynamics therefore exhibit a characteristic two-stage structure (not visible at the time scale in Figure 1, but shown in a zoom in shown in Figure S1 for artificial saliva at 50% RH). The largest change in osmotic pressure (Δ*Π*) occurred at 30% RH (162.5 MPa), followed by 50% RH (94.0 MPa) and 70% RH (49.4 MPa).

Across all RH conditions, both organisms remained stable while droplets were liquid and *Π*_droplet_ was low, as well as after efflorescence when *Π*_droplet_ had reached equilibrium. Inactivation occurred only during efflorescence, when rapid solute concentration increases produced a sharp rise in osmotic pressure. For *E. coli*, the magnitude of inactivation depended strongly on RH: it exceeded the detection limit (>5 log_10_ reduction) at 30% RH, was approximately 3.8 log_10_ at 50% RH, and approximately 1 log_10_ at 70% RH. *S. epidermidis* followed the same qualitative pattern but experienced less inactivation, with approximately 1.7 log_10_ decay at 30% RH, 1.5 log_10_ at 50% RH, and 0.7 log_10_ at 70% RH. In contrast to viability, the concentration of bacterial genome copies remained relatively constant throughout the experiment (Figure S2), indicating that the observed decay resulted from inactivation rather than adsorption of bacteria to the well plates. Neither *E. coli* nor *S. epidermidis* showed significant decay when held for 24 hours in bulk saturated NaCl solutions, where no rapid changes in osmotic pressure occur (Figure S3).

### Association between inactivation and rate of osmotic pressure change

Given that inactivation only occurred when *Π*_droplet_ increased (Fig. 1), we determined that the loss of viable titer is associated with the rate of change in osmotic pressure (Δ*Π/*Δ*t*). To this end, additional inactivation experiments were conducted over an RH range from 40 to 80 %, as well as in diluted and concentrated artificial saliva in order to see the effect of different initial osmotic pressures on inactivation (Figures S4, S5, S6; Table S1). Δ*Π/*Δ*t* at each RH was determined by ResAM, using a 5-minute window spanning efflorescence (4 minutes before and 1 minute after). The selected timescale also falls within the period over which *E. coli* cannot yet mount a full osmoadaptive response, which typically requires tens of minutes. Correlations were constructed across all artificial-saliva droplet experiments, relating the rate of osmotic pressure change around efflorescence to the corresponding reduction in bacterial titer. As shown in Table 1 and Figure 2, the decrease in viable titer following efflorescence was log-linearly related to Δ*Π/*Δ*t* for both bacteria.

**Table 1:**
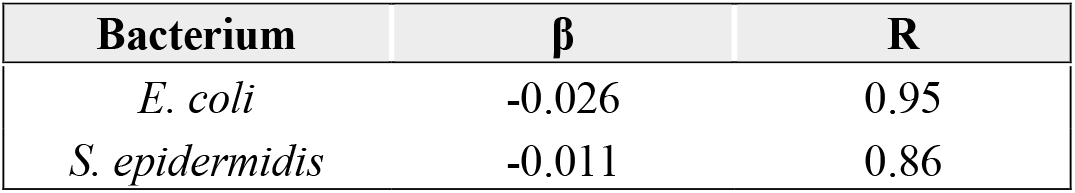
Log-linear regression coefficients for the relationship *log*_*10*_*(C/C*_*0*_*) = β(*Δ*Π/*Δ*t). β* is the slope associated with osmotic pressure change rate (MPa·5 min^−1^). *R* indicates the Pearson correlation coefficient.

| Bacterium | $\beta$ | R |
| --- | --- | --- |
| *E. coli* | -0.026 | 0.95 |
| *S. epidermidis* | -0.011 | 0.86 |

**Figure 2:**
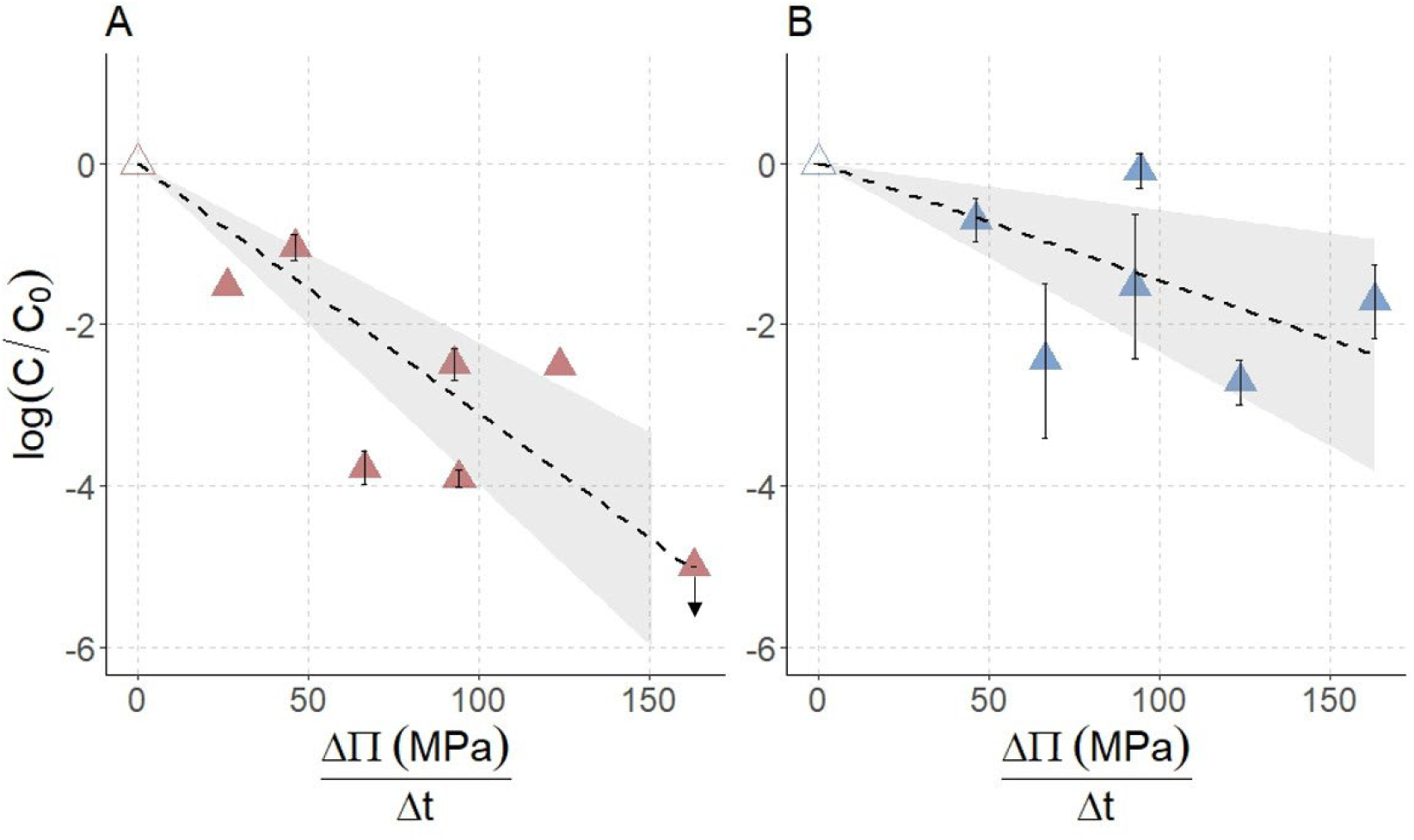
Correlation of the maximum change of osmotic pressure (Δ*Π/*Δ*t*) 4 minutes before and 1 minute after efflorescence and the inactivation of *E. coli* (A) and *S. epidermidis* (B) measured after efflorescence The error bars represent the standard deviation of all of the data points after efflorescence. The empty triangle represents experiments performed in bulk 5.4 M NaCl (Table S6). The dashed lines show the fits obtained by linear regression and the grey area indicates the associated 95 % confidence intervals. The arrow on the *E. coli* graph shows that the point was placed at the detection limit and could be lower (see Figure 1 Panel A).

To assess whether this relationship reflects a general physical mechanism rather than a medium-specific effect, independent droplet experiments were performed in phosphate-buffered saline (PBS) at 60% RH for *E. coli* and 40% RH for *S. epidermidis*. Using the *ΔΠ/Δt*–*inactivation* relationships derived from artificial saliva (Figure 2), predicted post-efflorescence titers in PBS were calculated via ResAM and compared with experimentally measured values. PBS was chosen because it is chemically distinct and simpler than artificial saliva, allowing us to test whether the rate of osmotic pressure change alone can predict bacterial survival.

As shown in Figure 3, predicted titers for both organisms closely tracked the experimental measurements in PBS. Model agreement with post-efflorescence data was evaluated by quantifying the fraction of observations falling within the 95% confidence interval and by comparing observed and predicted values using paired Wilcoxon signed-rank tests. For *E. coli*, 75% of observations fell within the confidence interval, and observed values did not differ significantly from predictions (*p = 0*.*20*), indicating good agreement. For *S. epidermidis*, 89% of observations fell within the confidence interval, but observed values differed significantly from predictions (*p = 0*.*0039*), indicating a systematic overestimation of inactivation despite the model capturing most of the observed variability.

**Figure 3:**
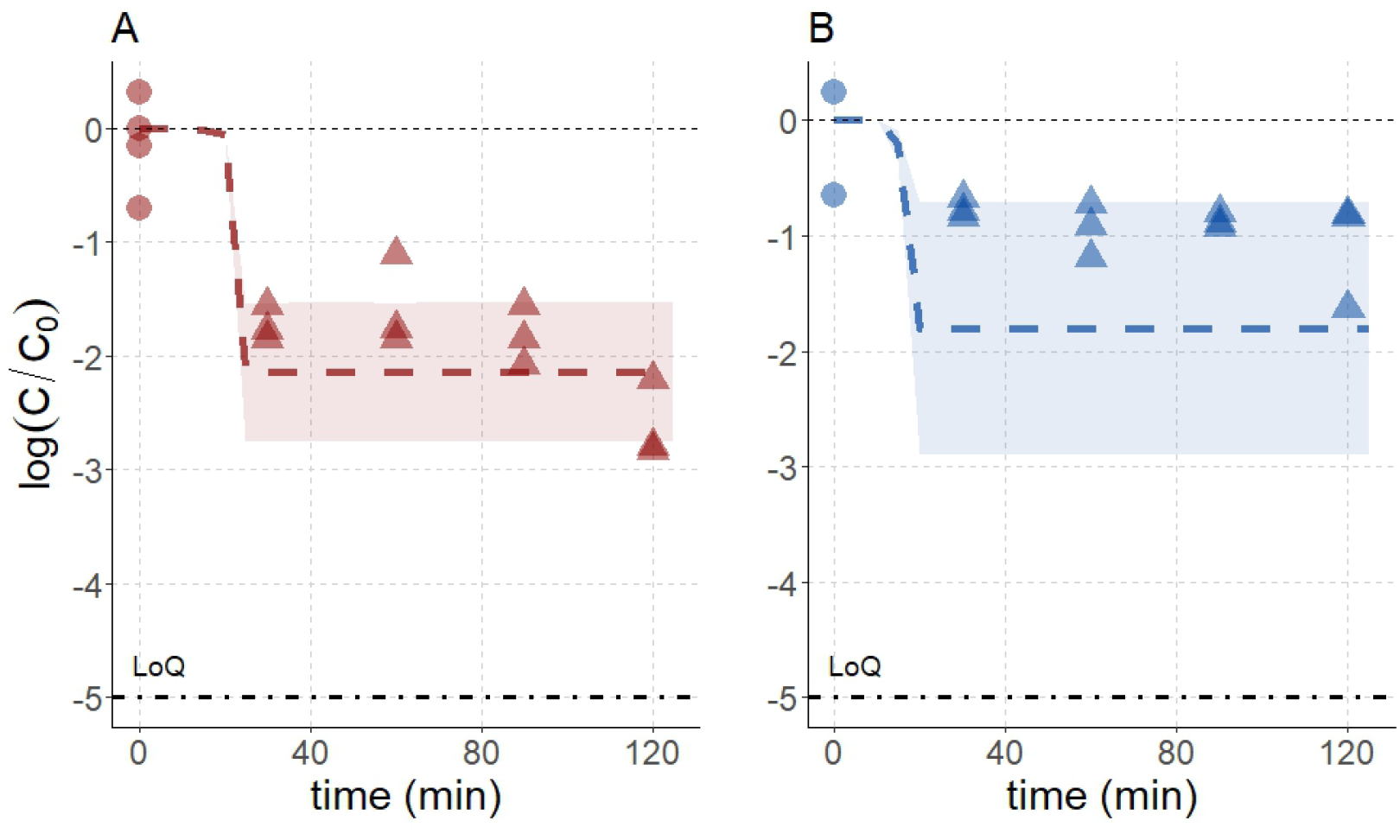
Bacterial titers of *E. coli* at 60% RH in PBS (A), and of *S. epidermidis* at 40% RH in PBS (B). Dashed lines represent model predictions. Data points indicate experimental measurements for *E. coli* (red) and *S. epidermidis* (blue), whereby circles represent liquid droplets and triangles represent effloresced droplets. The shaded regions represent the 95% confidence interval. Predictions were performed based on simulations of Δ*Π/*Δ*t* in ResAM, combined with the Δ*Π/*Δ*t*–inactivation relationships from artificial saliva in Figure 2.

Together, these results demonstrate that Δ*Π/*Δ*t*–inactivation relationships capture the timing and magnitude of bacterial inactivation in PBS, supporting the rate of osmotic pressure change at efflorescence as a broadly predictive, medium-independent determinant of bacterial survival.

### Hyperosmotic shock at efflorescence drives inactivation despite decreasing RH

Having established that bacterial inactivation scales with the magnitude of the osmotic pressure change around efflorescence, the inactivation dynamics of *E. coli* during droplet drying under decreasing RH conditions were next investigated. Because osmotic pressure increases as droplets dry but rises most sharply at efflorescence, varying the rate of RH decrease alters the osmotic pressure trajectory and thereby controls both the timing and magnitude of this critical osmotic shock.

Two droplet experiments were performed: one with a rapid decrease in RH from 90% to 30% and one with a gradual decrease (Figure S7). By comparing osmotic pressure, efflorescence timing, and post-efflorescence titers, it was assessed whether bacterial inactivation depends primarily on the abrupt osmotic pressure jump at efflorescence or on the slower osmotic pressure increase during drying.

Figure 4 shows that *E. coli* viability is governed by the rapid osmotic pressure increase immediately before efflorescence. Although osmotic pressure rises progressively even after efflorescence, the decay in *E. coli* titer is only minor, indicating that this gradual increase is not sufficient to impose strong osmotic shock on the bacteria.

**Figure 4:**
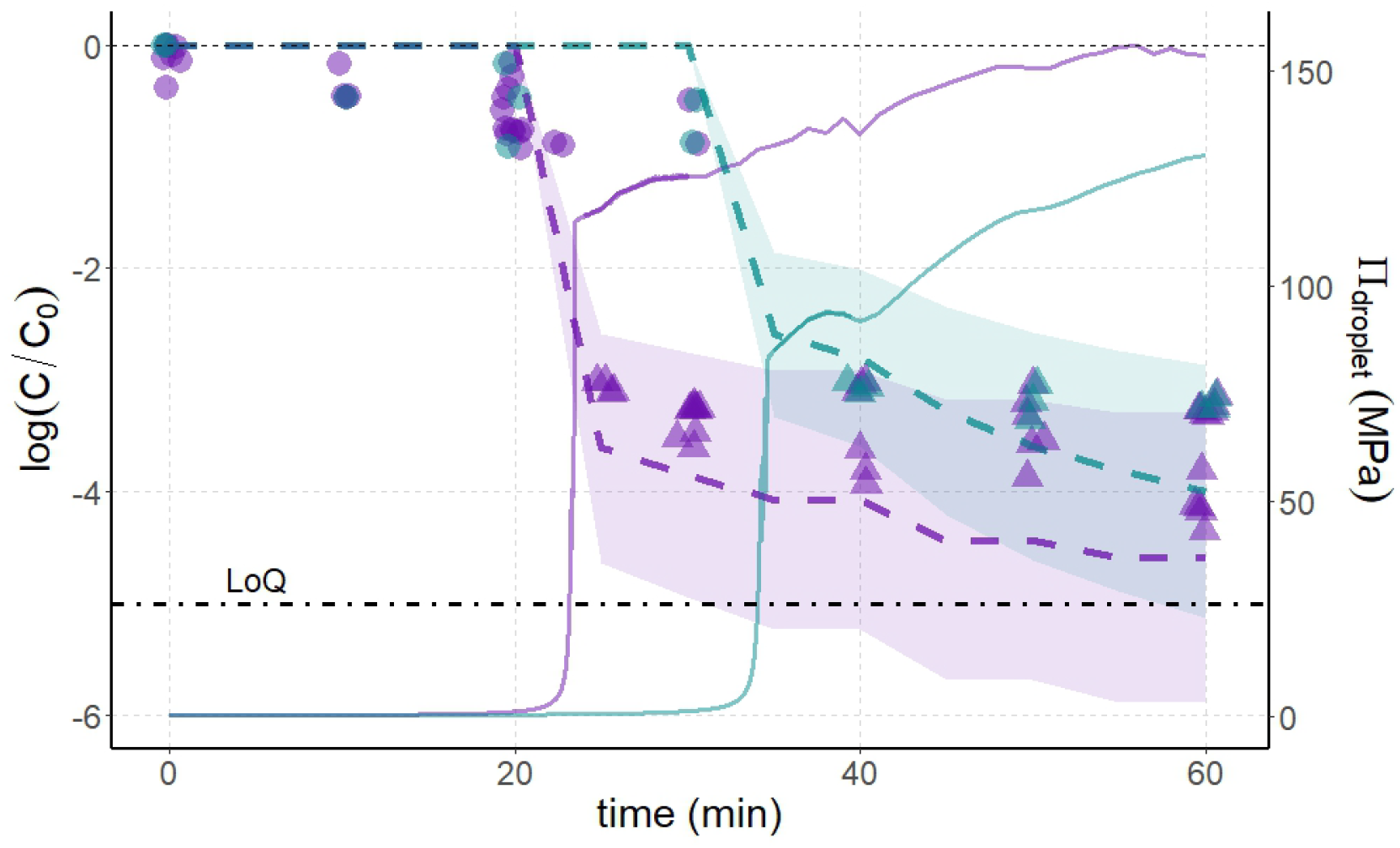
Fast relative humidity decrease (purple) and slow relative humidity (teal) with corresponding *E. coli* titers (circles represent liquid droplets and triangles represent effloresced droplets). The dashed line represents modeled *E. coli* from the model in Figure 2. The solid line represents modeled osmotic pressure in the two experiments from ResAM. The shaded regions are the 95% percent confidence intervals for the linear model.

In the rapid drying experiment (purple), efflorescence occurred at about 23 minutes corresponding to a Δ*Π/*Δ*t* of 118 MPa. In the slow drying experiment (teal), efflorescence occurred at about 34 minutes corresponding to a Δ*Π/*Δ*t* of 85.9 MPa. Inactivation was greater under rapid drying (3.8 log_10_) than under slow drying (3.2 log_10_), consistent with efflorescence occurring at a lower RH and producing a larger osmotic pressure jump than in the slower drying case. Using *the* Δ*Π*-inactivation relationship in Figure 2, the fast experiment has a predicted decay of 3.6 log_10_ while the slow experiment has a predicted decay of 2.8 log_10_.

To assess model agreement with the post-efflorescence data, the fraction of observed points falling within the 95% confidence interval of the model predictions was quantified, and observed and predicted values were compared using paired Wilcoxon signed-rank tests.

Under the fast-drying condition, 79.5% of post-efflorescence observations fell within the model confidence interval, and observed values did differ significantly from predictions (*p =* 5.45 · 10^-8^) indicating systematic overestimations of the decay. Under the slow-drying condition, all observations fell within the confidence interval, and observed values did not differ significantly from model predictions (*p = 0*.*075*), indicating good model agreement.

## Discussion

A quantitative relationship was established between the rate of change in osmotic pressure and the loss of bacterial viability of *E. coli* and *S. epidermidis* in artificial saliva droplets. Importantly, combining the empirically derived Δ*Π/*Δ*t–*inactivation relationship enabled prediction of both the timing of efflorescence during drying and the associated bacterial inactivation. The results indicate that bacterial survival is governed not by the final osmotic pressure reached, but by the rate of change in osmotic pressure right before efflorescence that exceeds bacterial adaptive capacity. Because this rate of change is strongly controlled by ambient relative humidity, small changes in indoor humidity can substantially alter microbial survival during droplet drying. By resolving the timing and magnitude of these short-lived events, the model provides a mechanistic framework linking the dynamics of droplet drying to the observed viability loss.

While both organisms exhibit increased inactivation with increasing rate of osmotic pressure change in droplets, *E. coli* is more susceptible to osmotic pressure changes compared to *S. epidermidis*. Inactivation was modest for both organisms at high relative humidity (70% RH), but diverged under typical indoor conditions: at 50% RH, *E. coli* exhibited ∼3.8 log_10_ inactivation compared to ∼1.5 log_10_ for *S. epidermidis*, with this disparity increasing further under drier conditions. This finding is consistent with prior reports on the relative sensitivities of these two organisms to desiccation and osmotic stress and implies that RH control is a more effective tool for inactivating Gram-negative than Gram-positive bacteria in IRPs^9^. The stronger susceptibility of *E. coli* relative to *S. epidermidis* supports the broader paradigm that Gram-negative bacteria are less tolerant of rapid changes in osmotic pressure than Gram-positive organisms^14^.

### Comparison to prior research

This study contextualizes prior work examining bacterial responses to osmotic and evaporative stress and extends these studies by quantitatively resolving the rapid osmotic transitions occurring at droplet efflorescence. Poirier et al. investigated *E. coli* responses to controlled hyperosmotic shocks by modulating water activity with glycerol and showed that viability depends strongly on the rate of osmotic change, with an optimal window in which cells can adapt^8^. Although the magnitude of osmotic pressure changes imposed in their system was comparable to those modeled in our droplets, their observed survival of *E. coli* was substantially higher. his likely reflects the protective effects of glycerol and other organic osmolytes, which can reduce membrane damage in bacteria ^20^. Their results highlight that *E. coli* can tolerate gradual osmotic increases but not rapid ones, supporting our conclusion that the abrupt Δ*Π* associated with droplet efflorescence exceeds the physiological tolerance of Gram-negative bacteria.

In addition to osmotic shock, other studies have identified oxidative stress as a potentially important inactivation mechanism in evaporating droplets. Liang et al. examined *E. coli* inactivation in artificial saliva droplets across RH gradients and separated the contributions of osmotic stress and thiocyanate (SCN^-^) toxicity ^21^. They found that osmotic stress dominated at low RH, while chemical toxicity became more influential as RH increased. Although their artificial saliva formulation differed from ours, their observation that faster evaporation intensifies osmotic stress complements our mechanistic interpretation. Our modeling adds to their framework by quantifying the sharp osmotic increase immediately before efflorescence, a transient event that cannot occur in bulk and was not captured in their analysis but directly corresponds to the period of maximum inactivation in our experiments.

Other work has proposed oxidative stress as a dominant inactivation mechanism in small, levitated aerosol droplets^22,23^. These aerosol studies use micrometer-scale droplets with high surface-to-volume ratios, which enhances interactions with reactive oxygen species (ROS) and produces stress pathways distinct from those in sessile droplets. This contrast helps explain why oxidative stress appears less relevant in our system: in addition to having possibly less ROS in our experiments, the lower surface to volume ratio of sessile droplets slows ROS accumulation, whereas water still equilibrates rapidly after efflorescence and generates a brief but extreme hyperosmotic shock which these aerosol studies cannot reproduce. This comparison underscores that dominant inactivation mechanisms shift with droplet size and evaporation dynamics.^21^

Taken together, these studies contextualize our results by showing that while bacteria can withstand gradual osmotic changes or experience additional stresses in aerosols, the rapid, efflorescence-associated osmotic surge unique to drying droplets imposes a level of stress that neither *E. coli* nor *S. epidermidis* can physiologically accommodate.

### Comparison to respiratory pathogens

Although *E. coli* and *S. epidermidis* serve as surrogates in this study, pathogenic Gram-positive respiratory bacteria often show strong desiccation tolerance: *Staphylococcus aureus* dried in PBS on polystyrene or stainless steel at 20–25% RH, ∼22 °C, exhibited <1 log_10_ loss after 7 days and only 1.5–2 log_10_ loss after 14–21 days^24^. Similarly, *Streptococcus pneumoniae* dried from Todd-Hewitt broth with yeast onto plastic surfaces at 20–30% RH, 23–25 °C, showed <1 log_10_ reduction after 24 h, with viable cells recovered after 4 days and 20–30% survival depending on serotype, indicating that Gram-positive respiratory pathogens generally retain substantial viability under low-RH drying conditions ^25^. Based on their shared Gram-positive physiology, many respiratory pathogens would be expected to display comparable resilience to our *S. epidermidis* surrogate under droplet-drying conditions.

Several nontuberculous mycobacteria, which are taxonomically Gram-positive Actinobacteria, exhibit substantial desiccation tolerance. Water-acclimated *Mycobacterium avium* and *M. chimaera* dried on filter paper at approximately 40% RH and 25 °C retained 28% and 34% viability after 21 days, with one *M. chimaera* strain showing 100% survival over the same period ^26^. When dried as biofilms on stainless steel under the same conditions,

*M. intracellulare* showed greater than 100% survival after 42 days, while *M. avium* and *M. abscessus* retained 18% and 14% survival, and *M. chelonae* survived up to 21 days. These results demonstrate that several respiratory-relevant bacteria with thick, complex cell envelopes can remain viable for extended periods at low relative humidity. Within this broader context, *E. coli* represents a highly sensitive, low-survival case, whereas *S. epidermidis* provides a more appropriate benchmark for the desiccation tolerance expected of clinically relevant respiratory bacteria in drying droplets.

In contrast to bacteria, previous work on influenza A virus in similar drying droplets reported continued inactivation after efflorescence, with a transition to slower, first-order decay rather than a plateau^4^. This difference reflects distinct inactivation mechanisms. For influenza virus, inactivation is driven by sustained exposure to high salt concentrations and associated damage to virion structure, allowing decay to continue after equilibration. In contrast, bacterial inactivation in this study is confined to the brief period of rapid osmotic pressure increase around efflorescence, after which viability remains stable. This suggests that bacterial inactivation is driven by acute osmotic shock rather than prolonged exposure to elevated salinity.

### Implications for airborne IRPs

This study focuses on sessile droplets, though bacterial inactivation in airborne IRPs is likely a larger point of concern. It is therefore important to relate the findings from this work to bacterial inactivation in smaller, airborne IRPs. Small IRPs (<10–20 μm) evaporate extremely rapidly, cool significantly, and have much higher surface-to-volume ratios than the millimeter-scale droplets used here ^27^. These features accelerate solute concentration and also promote accumulation of reactive oxygen species, producing stress pathways likely not relevant in larger droplets ^22^. Importantly, aerosols also undergo rapid pH changes during evaporation, driven by volatilization of CO2, partitioning of volatile acids and bases, and concentration of buffering species ^28,29^. Consequently, while aerosol inactivation reflects the combined influence of osmotic shock, oxidative stress, and rapid pH shifts, the much shorter evaporation timescales in airborne IRPs amplify the rate of osmotic pressure change to an even greater degree, suggesting that osmotic shock remains a key driver of bacterial loss in this regime ^28^.

### Limitations

This study has several limitations. Artificial saliva was used as the respiratory matrix, but mucin was excluded to enable more accurate modeling of osmotic pressure. Although mucin can alter droplet evaporation dynamics, previous work suggests it does not substantially affect bacterial survival^23^. Nevertheless, droplet drying dynamics in this study may differ from those of real saliva. Another study by Longest et al. shows that using pooled human saliva shows a greater rate of inactivation of Influenza A virus, likely due to the presence of additional chemical constituents, including proteins, enzymes, and reactive species, that are not present in artificial saliva ^30^.

Second, the bacterial species used in this study, *E. coli* and *S. epidermidis*, serve as model organisms rather than true respiratory pathogens. Although they represent key structural differences between Gram-negative and Gram-positive bacteria, clinically relevant respiratory pathogens may possess distinct survival strategies, surface characteristics, and stress-response kinetics. Therefore, extrapolation of these findings to pathogenic species should be made with caution until similar experiments are conducted directly on those organisms.

Third, the experiments were performed using 1 μL droplets, which are larger than most respiratory droplets generated during breathing, speaking, or coughing. Because droplet size strongly influences evaporation dynamics, solute concentration rates, and the timing of efflorescence, smaller droplets may experience different osmotic pressure trajectories than those observed here. Future work should investigate whether similar osmotic shock dynamics occur in droplets more representative of airborne respiratory particles.

Additionally, the Δ*Π/*Δ*t–*inactivation relationship used to relate bacterial inactivation to the rate of osmotic pressure change (Figure 2) was constrained to a 5-minute time window around efflorescence. This limited the range of osmotic pressure changes captured by the model. The constraint was particularly relevant for 10× artificial saliva, where osmotic pressure increased more gradually and therefore reached a lower value within the 5-minute window than in 1× or 0.1× artificial saliva (Figure 2). Extending the observation window to 10 minutes would likely reduce these differences, as the full change in osmotic pressure would be more completely captured over that longer timeframe (see Figure S4 for the evolution of osmotic pressure of a 10x Artificial Saliva Droplet).

The physicochemical simulations were also subject to limitations associated with the ResAM framework. In particular, the model relies on assumptions regarding droplet composition, thermodynamic equilibrium, and homogeneous solute distribution during evaporation. These simplifications may not fully capture microscale concentration gradients, phase transitions, or spatial heterogeneity that can arise within drying respiratory droplets. Additional assumptions and parameter uncertainties associated with the model are summarized in the Supplementary Information.

Taken together, this droplet-based system offers a controlled framework for quantifying osmotic shock and its effects on bacterial survival. However, it does not fully replicate the complex physicochemical environment of airborne respiratory aerosols. Future work extending this approach to airborne infectious respiratory particles will be necessary to capture the additional stress factors that influence microbial viability under real-world transmission conditions.

## Materials and Methods

### Osmotic pressure modelling

Each droplet was modeled using the Respiratory Aerosol Model (ResAM), a spherical shell diffusion model that simulates the evolution of physicochemical conditions within drying respiratory droplets. The model resolves radial concentration gradients and water activity across concentric shells as evaporation proceeds. The model was initialized using the measured environmental conditions inside the chamber, including relative humidity, and temperature together with the initial droplet composition and the experimentally measured droplet size evolution obtained from video recordings.

The original version of ResAM is fully described in Luo. et al. ^28^, which was adapted for virus-containing droplets seen in Schaub et al.^4^. For droplets containing *E. coli* and *S. epidermidis*, the model was further adapted to account for bacteria changing the droplet morphology and increasing the rate of evaporation ^31^. The adaptations are described in the SI (Method S1). Water activity values simulated by ResAM were used to calculate the osmotic pressure within each droplet shell according to the van’t Hoff equation ^9^:

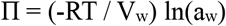

Where *Π* is the osmotic pressure in MPa, R is the universal gas constant (8.314 J mol^-1^ K^-1^), *T* is temperature in Kelvin (298 K), *V*_*w*_ is the molar volume of water at 298K (18.015 × 10^-6^ m^3^), and *a*_*w*_ is the water activity output of ResAM. This formulation defines osmotic pressure relative to pure water (*a*_*w*_ = 1). The osmotic pressure was averaged across shells to generate the lineplot visualizations. ResAM simulations were executed on a high-performance computing cluster, and the resulting shell-resolved outputs were processed and analyzed in R (version 2023.09.1).

### Droplet Experiments

Droplet experiments were performed in an environmental chamber (Electro-tech Systems 5332) with controls for relative humidity and temperature. Prior to the experiment and during bacterial resuspension, the chamber was set to the desired relative humidity and temperature. Once the chamber was ready, 1 μL droplets were deposited into a 96-well plate with non-binding surfaces (Greiner Bio-One, 655901), with one droplet per well. Droplets were left to dry over the course of 60 min for 30% (only for *E. coli*) and 120 min for other relative humidities and were periodically collected at each time point, with three droplets deposited per time point. The first three deposited droplets were collected immediately and were considered the t_0_ time point.

Droplets were collected by depositing 300 μL of PBS into the well. The droplets were then resuspended by pipetting up and down with a 200 μL pipette ten times, scratching the bottom of the well with the pipette tip 5 times, and then repeating those two steps. Each sample was aliquoted into two 150-μL samples, one placed into a separate 96-well plate (Greiner) for an infectivity assay and the other into an Eppendorf tube for dPCR analysis. It was visually inspected and noted if the droplet was crystallized or not. In each experiment, a control was included by sampling 1 μL of the experimental solution directly from the Eppendorf at both the start and conclusion of the experiment and kept inside the chamber to maintain the same conditions as the droplet experiment. See Table S2 for experimental parameters.

Experiments at constant saturated NaCl were conducted in bulk solutions. See Supporting Information (Method S5) for details.

### Determination of ΔΠ/Δt–inactivation relationship

To quantify the rate of osmotic pressure change, the efflorescence time was first identified for each droplet. A 5-min window spanning this event (4 min before and 1 min after efflorescence) was then extracted. This interval captures the complete efflorescence transition across all experiments while remaining short relative to the droplet sampling timescale.

Within this window, the change in osmotic pressure (Δ*Π/*Δ*t*) was calculated from the ResAM output and related to the experimentally observed inactivation using linear regression. The selected timescale also falls within the period over which *E. coli* cannot yet mount an osmoadaptive response, which typically requires tens of minutes. We therefore assume that *Π*_bacteria_ does not change over the 5 min window considered; the change in turgor pressure P_turgor_ before and after efflorescence, (*Π*_bacteria_ *-Π*_droplet, before_) - (*Π*_bacteria_ *-Π*_droplet, after_), then corresponds to Δ*Π*_droplet_ *= Π*_drople*t*, after_ *-Π*_droplet, before_, here simply termed Δ*Π*.

Post-efflorescence bacterial titers were averaged and linked to the corresponding osmotic pressure change, and the resulting fit was used to predict bacterial inactivation at efflorescence. The point at the origin represents the osmotic pressure difference between PBS, in which the bacteria were initially suspended, and a 5.4 M NaCl solution (see Methods S2 and S3).

### Prediction of bacterial titers

To predict bacterial titers, Δ*Π/*Δ*t* was calculated from the ResAM output by determining the slope over consecutive 5 min intervals. These values were then used together with the experimentally derived Δ*Π/*Δ*t–*inactivation relationship to estimate bacterial decay at each interval. Cumulative bacterial inactivation was calculated by summing the predicted decay across successive intervals. The model assumes that inactivation is primarily associated with efflorescence; therefore, predicted bacterial titers remain constant thereafter unless additional osmotic changes occur. Post-efflorescence, decay was only applied when the change in osmotic pressure exceeded 5 MPa; otherwise, titers were assumed to remain unchanged.

### Organism Propagation, Preparation and Enumeration

Detailed methods for organism propagation, preparation, and enumeration are provided in the Supporting Information (Method S4)

### DNA extractions and dPCR quantification

Detailed protocols for DNA extraction and dPCR quantification are provided in the Supporting Information (Method S5).

### Determination of Crystallization Time

Methods used to determine crystallization time are described in the Supporting Information. (Method S6).

### Changing relative humidity experiments

Detailed experimental procedures for changing relative humidity experiments are described in the Supporting Information (Method S7).

### Statistical Analysis

All statistical analyses were performed in R (version 2023.09.1). Linear regression was used to quantify relationships between osmotic pressure change and bacterial inactivation.

Differences between predicted and observed titers were evaluated using paired Wilcoxon signed-rank tests. Efflorescence time distributions were fitted using the *fitdistrplus* package in R. Significance was assessed at α = 0.05. Error bars represent the standard deviation of biological replicates.

## Supporting Information

Supporting methods, supporting figures, and supporting tables are available in the supporting information.

## Data availability

Experimental data are available in the public repository Zenodo under the following link: 10.5281/zenodo.19083698.

All code is available at: https://github.com/medinataylor/dropletpaper

## Acknowledgements

The author thanks Dr. Priya Ramakrishna for assisting with osmotic pressure explanations. This work was supported by an internal grant (ENAC flagship grant ICARUS) and by the Swiss National Science Foundation grant no. IC00I0-228106.

## 8. Author Information

TM contributed to Conceptualization, Methodology, Software, Validation, Formal Analysis, Investigation, Data Curation, Writing – Original Draft, Writing – Review & Editing, and Visualization. BL contributed to Software, Validation, Formal Analysis, and Writing – Original Draft (Supporting Information only). TP contributed to Validation and Writing – Review & Editing. HKW contributed to Investigation. TK contributed to Conceptualization, Methodology, Validation, Writing – Review & Editing, Supervision, and Funding Acquisition.

