## Supplemental methods, figures and tables for "Rate of osmotic pressure change in drying saliva microdroplets drives inactivation of surrogate respiratory bacteria"

**This file includes:**

20 pages

Supporting Methods S1-S7

Figures S1 – S9

Tables S1 – S7

### Supplemental Methods

#### Method S1 - ResAM Model Adaptations

The full ResAM model for airborne particles is described in Luo et al. (2023)[1]. The model was subsequently extended by Schaub et al. (2024)[2] to account for droplets deposited on a substrate. In the present work, droplet flattening on a substrate was represented by incorporating an inert central core, the size of which was determined from experimental image analysis. The remaining liquid volume was assumed to be distributed as a hemisphere. To maintain geometric consistency, the measured image area was matched to the surface area of a hemisphere with a mean radius defined as the average of the inner and outer radii. The core size was treated as time-dependent and iteratively determined from the experimentally observed image area together with the calculated droplet volume.

Compared to the version presented by Luo et al. (2023), two major improvements were implemented:

1. **Multi-shell solidification**
   Solid formation was enabled in all shells of the droplet rather than being restricted to the center.
2. **Improved diffusion treatment**
   Experimentally realistic ion diffusion coefficients were incorporated for crystal growth calculations. This replaced the previous parameterized enhancement-factor approach used by Luo et al. (2023), which constrained crystallization to the droplet center.

In addition, the chemical system was substantially expanded to include a broader range of aqueous and solid-phase species.

### Chemical Species Included in the Model

The model included water and dissociated ions ($H_2O$, $H^+$, $OH^-$); two generic organic species with different molar masses and molar volumes; cations ($Na^+$, $K^+$, $Ca^{2+}$, $Mg^{2+}$); anions ($Cl^-$, $NO_3^-$); carbon species ($H_2CO_3$, $HCO_3^-$, $CO_3^{2-}$); ammonia species ($NH_3$, $NH_4^+$); acetic acid species ($CH_3COOH$, $CH_3COO^-$); phosphate species ($H_3PO_4$, $H_2PO_4^-$, $HPO_4^{2-}$, $PO_4^{3-}$); sulfate species ($HSO_4^-$, $SO_4^{2-}$); lactic acid species ($CH_3CH(OH)COOH$, $CH_3CH(OH)COO^-$); carbonic anhydrase enzyme; and additional acid species (default: oxalic acid) together with associated ions up to triple charge.

### Gas-Phase Species

The following gas-phase species were included in the model: $H_2O$, HCl, $CO_2$, $HNO_3$, $NH_3$, $CH_3COOH$, lactic acid, and an additional acid species (default: oxalic acid).

For this study, air chemistry interactions were simplified by assuming exchange with $CO_2$ only at a concentration of 400 ppm.

### Solid Phases Considered for Efflorescence

The following crystalline solids were included in the efflorescence calculations: NaCl, KCl, $CaCO_3$, $MgCO_3$, $CaHPO_4$, $NaH_2PO_4$, $Na_2HPO_4 \cdot 7H_2O$, $NH_4HC_2O_4 \cdot 0.5H_2O$, $(NH_4)_2C_2O_4$, $NaHC_2O_4$, $Na_2C_2O_4$, and $H_2C_2O_4 \cdot 7H_2O$.

Note that not all crystalline phases listed above were used in the present study. See Table S3 for what species were used in the model inputs.

### ResAM Model Inputs

Artificial saliva composition was based on the formulation provided by Pickering Laboratories (Table S3). Molality values used as model inputs were calculated from the reported composition. For PBS simulations, molality and molarity were assumed to be equivalent; therefore, the concentrations listed in Table S1 were used directly as model inputs. The initial droplet diameter used as input to ResAM was determined from videography measurements of the deposited droplets and was $7.82 \times 10^{-2}$ cm.

At 30% RH, ResAM calculations terminated following efflorescence. Consequently, osmotic pressure values beyond efflorescence were manually extended using the equation presented in the main text with a water activity value corresponding to equilibrium at 30% RH ($a_w$ = 0.3). The same approach was applied to the 50% RH simulations, where calculations terminated after 100 min and were extended to 120 min.

### Method S2 - Stability of Bacteria in Saturated Sodium Chloride Solution

To evaluate the effect of salt exposure alone on bacterial survival, 20 µL of either *E. coli* or *S. epidermidis* suspended in PBS was added to 180 µL of saturated NaCl solution (5.4 M) in technical triplicate at room temperature. Samples were collected immediately (time 0) and after 24 h. As shown in Figure S3, *S. epidermidis* remained stable over the 24 h exposure period, whereas *E. coli* exhibited approximately 1-log inactivation.

**Method S3 - Bulk Osmotic Pressure Calculations**

The osmotic pressure of the bulk solutions was calculated using the ideal solution osmotic pressure formula, with a correction factor for nonideality (ϕ) for the saturated NaCl solution.

$$\Pi = i\phi CRT$$

$\Pi$ is the osmotic pressure, $i$ is the Van't Hoff Factor, $C$ is the molar concentration, $R$ is the universal gas constant, and $T$ is the temperature. The empty triangle in Figure 2 was calculated using the difference of 5.4M NaCl and PBS, with a value of **33.5 MPa**. See Table S1 for osmotic pressure calculations.

**Method S4 - Organism Propagation and Purification**

Experiments were conducted using *Escherichia coli* (DSM-5695) and *Staphylococcus epidermidis* (DSM-1798). For *E. coli*, 10 mL Luria-Bertrani Broth (LB Broth) (Invitrogen 12795027) was added to a 15 mL falcon tube and 100 µL of *E. coli* stock was inoculated and then incubated in 37°C overnight with shaking at 200 RPM. Before each droplet experiment, the *E. coli* was vortexed and 1 mL was aliquoted into a 1.5 mL Eppendorf tube and centrifuged for four minutes at 10’000 x g. The supernatant was discarded and the pellet was resuspended in one mL of Artificial Saliva (Pickering Laboratories). For *S. epidermidis*, 10 mL of yeast-enhanced tryptic soy broth (TSB, Millipore 102512952 with 3g of yeast extract added per liter) was added to a 15 mL falcon tube and 100 µL of *S. epidermidis* was inoculated and then incubated in 37°C overnight. Resuspension into artificial saliva follows the same protocol as *E. coli*.

Live bacteria were quantified by plating 10-fold serial dilutions onto organism-specific agar media. *E. coli* was enumerated on LB agar (15% w/v agar), while *S. epidermidis* was enumerated on tryptic soy agar (TSA, Fluka Analytical 22091) supplemented with yeast extract. Plates were incubated at 37 °C for 18–24 h, after which colonies were counted and concentrations were reported as colony-forming units (CFU) on the order of $10^8$ CFU/mL.

**Method S5 - DNA Extraction and Digital PCR Quantification**

DNA from *E. coli* and *S. epidermidis* was independently extracted using the Maxwell RSC Instrument (Promega) together with the Maxwell RSC Whole Blood DNA Kit (AS1400) according to the manufacturer's instructions. DNA was eluted in 60 µL of supplied elution buffer and stored at −20 °C until quantification.

Extracted DNA was quantified using digital PCR (dPCR) with either the QIAcuity 16S *E. coli* Kit (Qiagen, cat. no. 18000-16) or the QIAcuity 16S *S. aureus* Kit (Qiagen, cat. no. 250207). All reactions were loaded into QIAcuity Nanoplate 8.5k 96-well plates (Qiagen, cat. no. 250021) and run on the QIAcuity One Digital PCR System (Qiagen, 5-plex device, cat. no. 911255).

The thermal cycling protocol consisted of enzyme activation at 95 °C for 2 min followed by 40 cycles of 95 °C for 15 s and 58 °C for 1 min. All dPCR runs included both positive controls (Qiagen, cat. no. 338135) and negative controls consisting of RNase-free water. To account for variations in recovery efficiency, CFU measurements were normalized to the number of genome copies detected at time 0, when droplets

**Method S6 - Videography and Efflorescence Determination**

Droplet evaporation and crystallization behavior were monitored using videography. Images were acquired using a Sony IMX477R camera connected to a Raspberry Pi 4 computer (Model B Rev. 1.4) powered by a Varta power bank. The camera was positioned 15 cm above the 96-well plate and captured one image every 16 s.

Droplet area and diameter were quantified using ImageJ (version 1.53t). A single well of the 96-well plate was used as the spatial reference for image calibration. Efflorescence times were determined by manually inspecting image sequences frame-by-frame and recording the onset and completion of crystallization events.

To determine efflorescence times in the presence of bacteria, droplets containing *E. coli* or *S. epidermidis* were monitored across multiple RH conditions. At 30% RH, *S. epidermidis* droplets effloresced earlier than *E. coli*, whereas both species exhibited similar efflorescence times at 50% RH (Figure S8, Table S2). At 70% RH, crystallization occurred during the experiment but could not be visually resolved due to limited contrast (Figure S9).

Consequently, ResAM was used to estimate efflorescence times across all RH conditions. Model-predicted values fell within the experimentally observed range for *E. coli* and were consistent with observations for *S. epidermidis* (Table S4).

### Method S7 - Relative Humidity Change Experiments

To investigate the effect of changing relative humidity (RH) on droplet behavior, two RH-transition protocols were performed.

For the fast RH-decrease experiment, the environmental chamber was initially equilibrated to 90% RH. Following droplet deposition, the chamber setpoint was switched to 30% RH, allowing the humidity to decrease naturally over the course of the experiment. Droplets were sampled at 0, 10, 20, 30, 40, 50, and 60 min, with additional sampling points at 22.5 and 25 min to capture the efflorescence transition.

For the slow RH-decrease experiment, the chamber was likewise initialized at 90% RH; however, the RH setpoint was reduced stepwise by 10% every 10 min. For ResAM simulations, the RH input for the fast-change condition was taken as the average RH measured across the two replicate fast-change experiments. See Figure S7 for the relative humidity change plot.

### Supplementary Figures

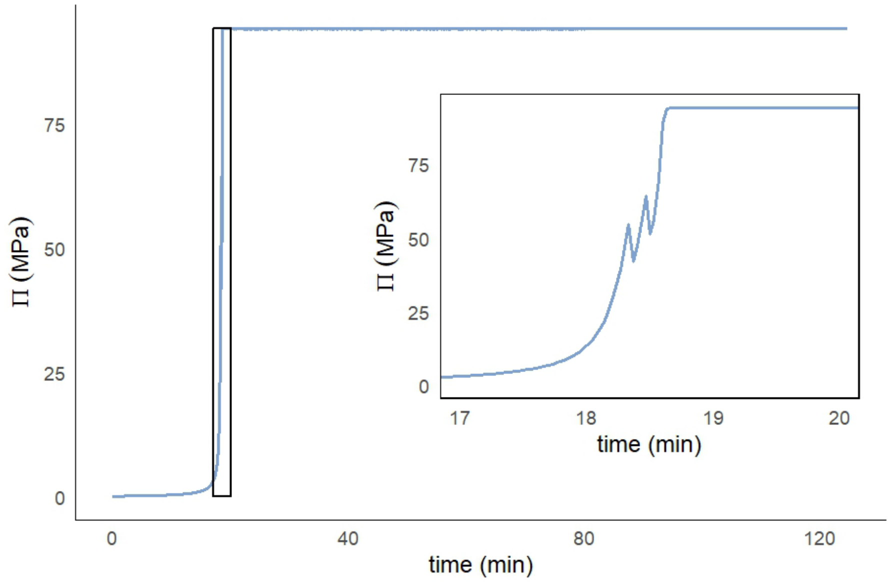


**Figure S1:** Zoomed-in osmotic pressure curve from Figure 1 for 1x Artificial Saliva at 50%.

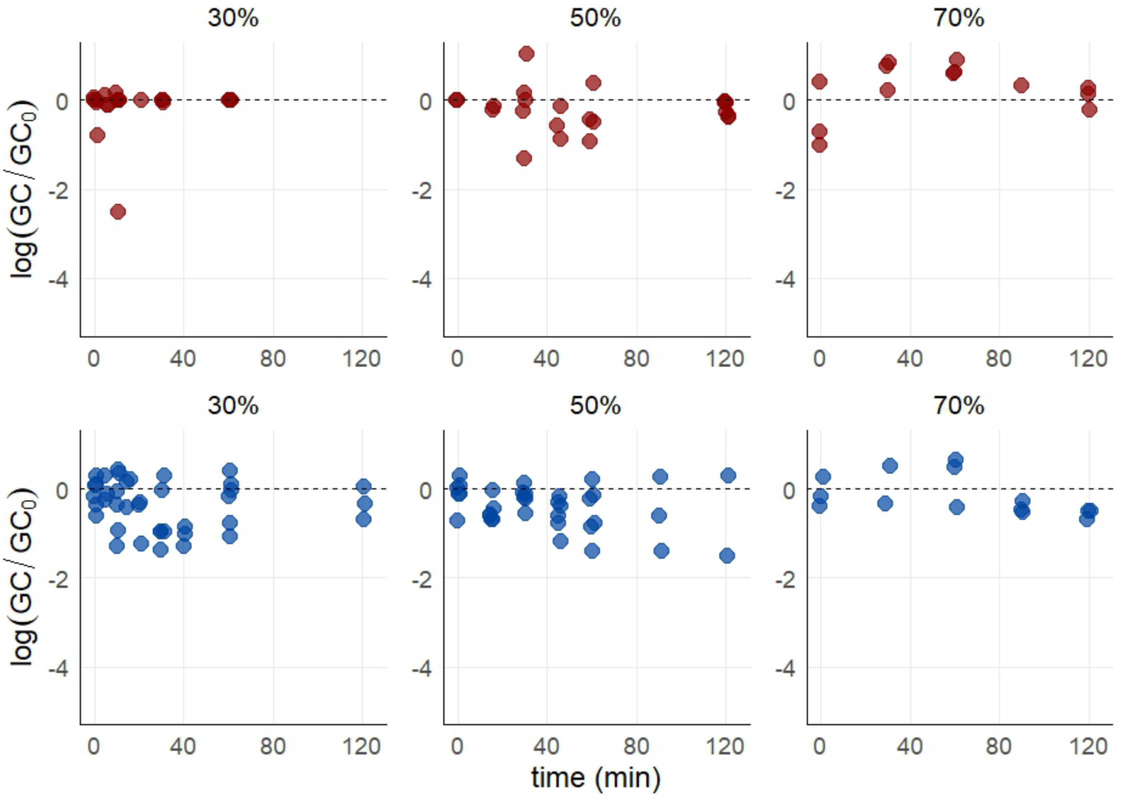


**Figure S2:** Genome copies (GC) normalized to time 0 for Figure 1. *E. coli* droplets are in red and *S. epidermidis* droplets are in blue.

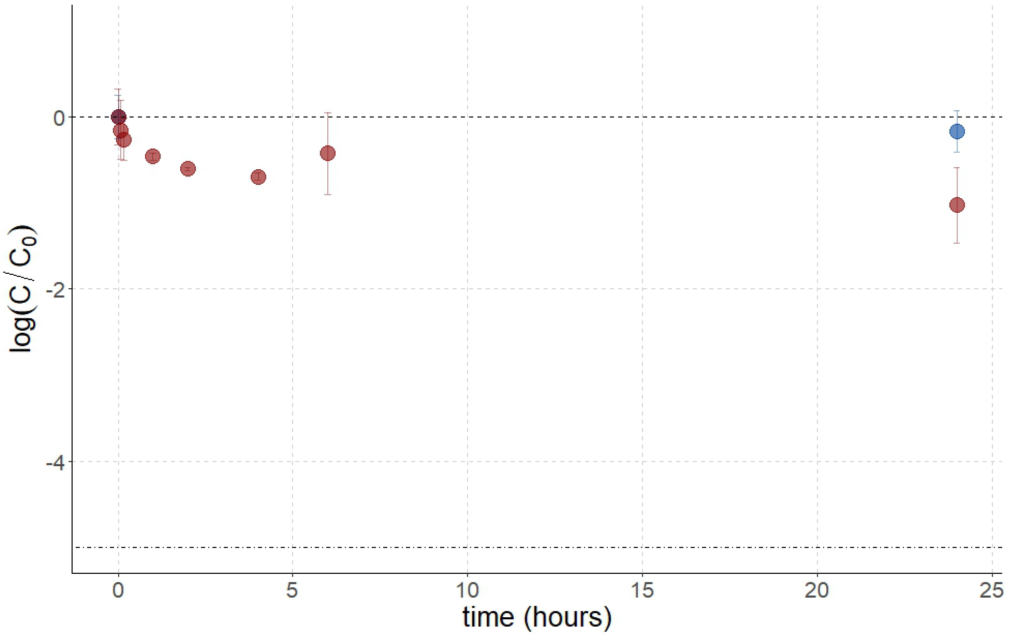


**Figure S3:** Stability of *E. coli* (red) and *S. epidermidis* (blue) in bulk 5.4M NaCl solution

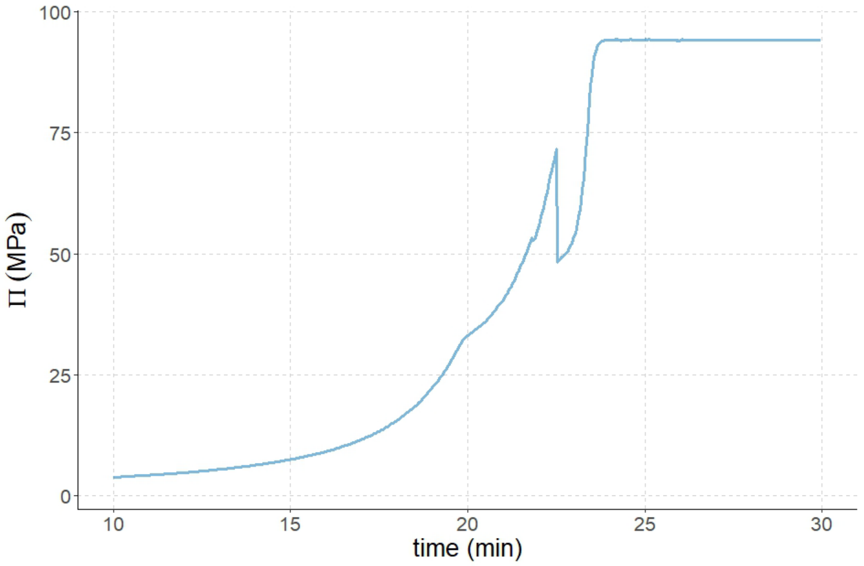


**Figure S4:** 10x Artificial Saliva osmotic pressure curve zoomed in from 10-30 minutes.

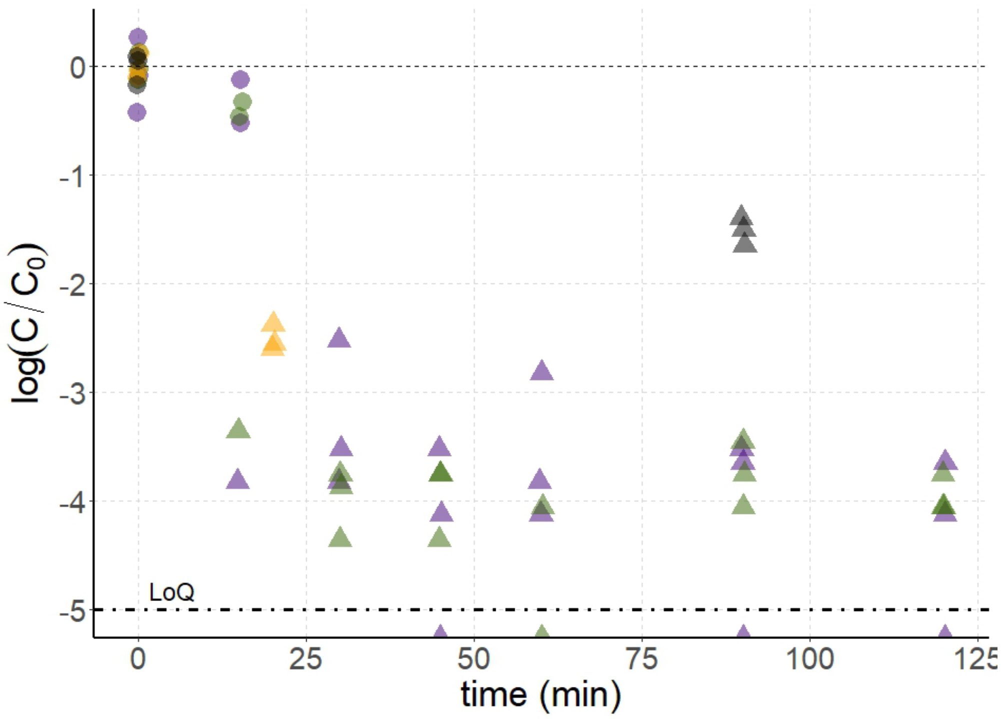


**Figure S5:** *E. coli* time series for 10x Artificial Saliva at 50% (purple), 0.1x Artificial Saliva at 50% (green), 40% (orange), and 80% (black).

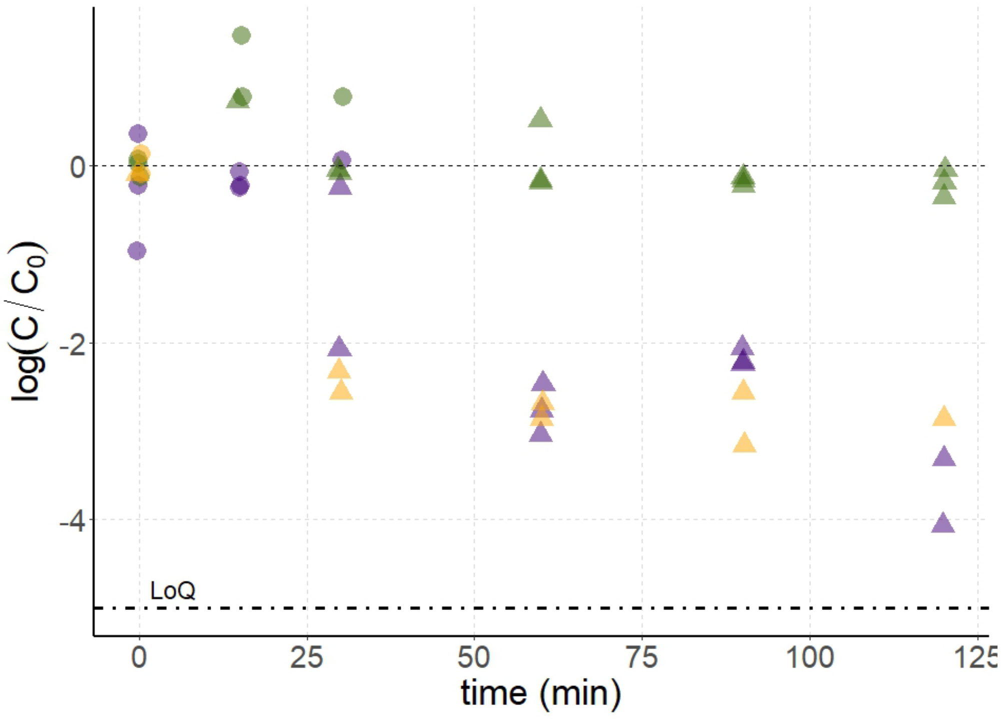


**Figure S6:** *S. epidermidis* time series 10x Artificial Saliva at 50% (purple), 0.1x Artificial Saliva at 50% (green), and 40% (orange)

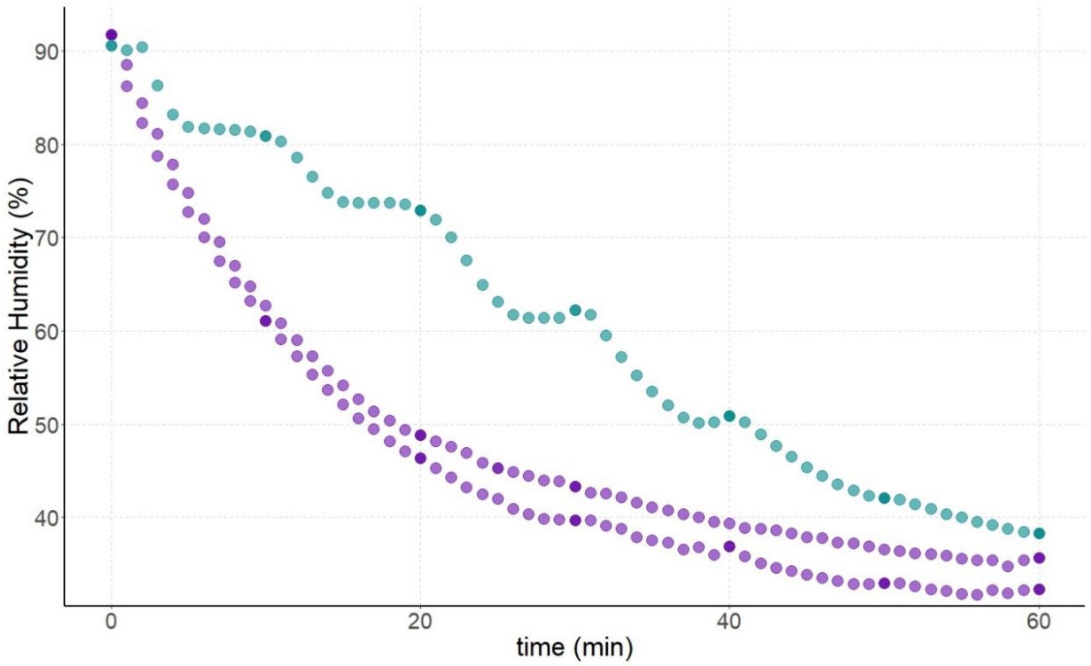


**Figure S7:** Relative Humidity change for the fast RH change experiments (purple) and the slow relative humidity change (teal) corresponding to Figure 4 in the main text.

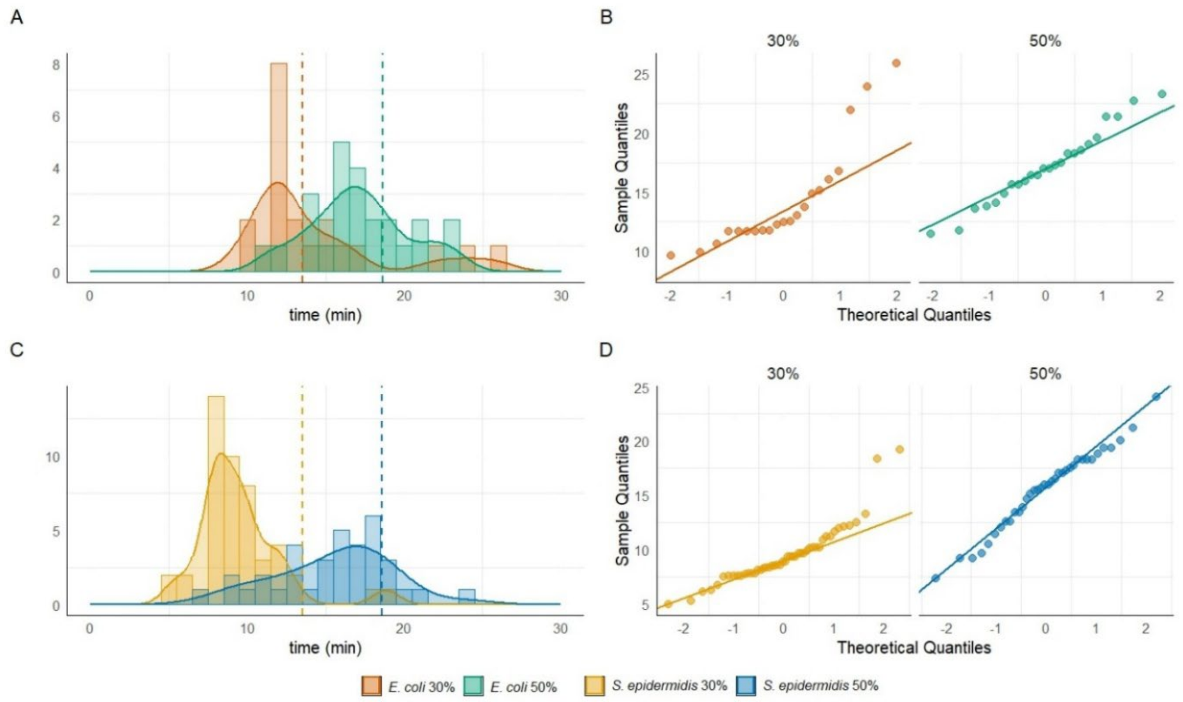


**Figure S8:** Panel A shows the distributions of droplet efflorescence times for *E. coli* at 30% RH (orange, n = 21) and 50% RH (green, n = 24), and panel B shows the corresponding Q–Q plots used to assess normality. Panel C shows the distributions of droplet efflorescence times for *S. epidermidis* at 30% RH (yellow, n = 48) and 50% RH (blue, n = 36), and panel D shows the corresponding Q–Q plots used to assess normality. Dashed vertical lines in the distribution plots indicate the model-predicted efflorescence times (13.5 min at 30% RH and 18.6 min at 50% RH).

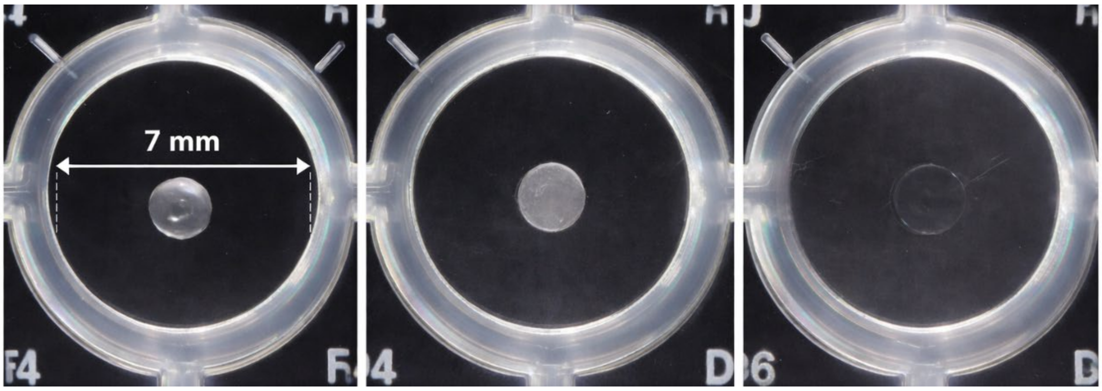


**Figure S9:** The photo on the left shows a liquid droplet, the photo in the middle shows an effloresced droplet, and the photo on the right shows a droplet at 70% that does not have a clear crystallization.

### Supplementary Tables

**Table S1:** Osmotic Pressure Calculations

| **Chemical** | **C (mol·L$^{-1}$)** | ***i*** | **R (L·MPa·mol$^{-1}$·K$^{-1}$)** | **T (K)** | **ϕ** | **Π (MPa)** |
|---|---|---|---|---|---|---|
| ***5.4M NaCl*** | | | | | | |
| NaCl | 5.4 | 2 | 0.008314 | 298 | 1.28[3] | **34.3** |
| ***PBS*** | | | | | | |
| NaCl | 0.137 | 2 | 0.008314 | 298 | 1.00 | 0.679 |
| KCl | 0.00270 | 2 | | | | 0.0134 |
| $Na_2HPO_4$ | 0.0100 | 3 | | | | 0.0743 |
| $KH_2PO_4$ | 0.00180 | 2 | | | | 0.00892 |
| Total PBS | | | | | | **0.775** |

**Table S2:** Summary of RH and Temperature Conditions by RH (μ ± σ)

| | ***E. coli*** | | ***S. epidermidis*** | |
|---|---|---|---|---|
| **Media** | **RH** | **Temperature ºC** | **RH** | **Temperature ºC** |
| 1x AS | 31.22 ± 1.46 | 22.56 ± 0.11 | 30.96 ± 0.71 | 23.95 ± 0.86 |
| | 42.56 ± 0.80 | 22.08 ± 0.16 | 42.95 ± 0.77 | 23.06 ± 0.76 |
| | 51.00 ± 1.45 | 22.62 ± 0.11 | 50.72 ± 0.23 | 24.55 ± 0.43 |
| | 69.81 ± 3.80 | 21.83 ± 0.85 | 70.43 ± 1.01 | 24.66 ± 0.46 |
| | 84.43 ± 1.50 | 23.52 ± 0.34 | | |
| 10x AS | 50.42 ± 4.43 | 21.09 ± 0.83 | 50.87 ± 0.72 | 24.85 ± 0.86 |
| 0.1x AS | | | | |
| PBS | 60.10 ± 1.20 | 22.85 ± 0.40 | 42.95 ± 0.77 | 23.06 ± 0.76 |

**Table S3:** Pickering Labs Artificial Saliva Composition

| **Component** | **Chemical Formula** | **Molar mass (g/mol)** | **Concentration (g/L)** | **Molality (mol / kg)** |
|---|---|---|---|---|
| Sodium chloride | $NaCl$ | 58.44 | 0.88 | 0.015 |
| Monopotassium phosphate | $KH_2PO_4$ | 136.09 | 0.21 | 0.0016 |
| Potassium chloride | $KCl$ | 74.55 | 1.04 | 0.014 |
| Sodium bicarbonate | $NaHCO_3$ | 84.01 | 0.42 | 0.0050 |
| Calcium chloride monohydrate | $CaCl_2 * H_2O$ | 129.00 | 0.13 | 0.0010 |
| Ammonium chloride | $NH_4Cl$ | 53.49 | 0.11 | 0.0021 |
| Dipotassium phosphate | $K_2HPO_4$ | 174.18 | 0.43 | 0.0025 |
| Potassium thiocyanate | $KSCN$ | 97.18 | 0.19 | 0.0020 |
| Magnesium chloride heptahydrate | $MgCl_2 * 7H_2O$ | 221.32 | 0.04 | 0.00018 |
| Urea | $CH_4N_2O$ | 60.06 | 0.12 | 0.0020 |

**Table S4:** Observed versus model droplet efflorescence times

| RH (%) | Bacterium | Observed Efflorescence Time (min) | Model Efflorescence Time (min) |
|---|---|---|---|
| 30 | *E. coli* | 15.6 ± 3.7 | 13.5 |
| | *S. epidermidis* | 9.5 ± 2.7 | |
| 50 | *E. coli* | 17.1 ± 3.1 | 18.6 |
| | *S. epidermidis* | 14.2 ± 4.5 | |
| 70 | *E. coli* | *not observable* | 30.9 |
| | *S. epidermidis* | *not observable* | |

**Table S5:** *E. coli* dPCR reaction mix

| ***E. coli*** **Assay** | **Reaction mix for 1 sample (1 µL)** |
|---|---|
| QIAcuity OneStep Advanced Probe – Mastermix (4x) | 3 |
| Microbial DNA Assay *E. coli* 20x (DMA00140-F) | 0.6 |
| GC enhancer | 1.5 |
| Ultrapure $H_2O$ | 2.9 |
| Sample | 4 |
| Final volume | 12 |

**Table S6:** *S. epidermidis* dPCR Reaction Mix

| ***S. epidermidis*** **Assay** | **Reaction mix for 1 sample (1 µL)** |
|---|---|
| QIAcuity OneStep Advanced Probe – Mastermix (4x) | 3 |
| Microbial DNA Assay *S. epidermidis* 20x (DMA00304-F) | 0.6 |
| GC enhancer | 1.5 |
| Ultrapure $H_2O$ | 2.9 |
| Sample | 4 |
| Final volume | 12 |

**Table S7:** dPCR protocol for *E. coli* and *S. epidermidis*

| **Step** | | **Temperature** | **Time** |
|---|---|---|---|
| PCR Initial Heat Inactivation | | 95ºC | 2 min |
| Denaturation | 40x | 95ºC | 15 s |
| Combined annealing / extension | | 58ºC | 1 min |
